# Synergistic anti-tumor activity of EGFR inhibition and C/EBPβ antagonism in GBM

**DOI:** 10.64898/2026.04.17.719281

**Authors:** Julia Diehl, Claudio Scuoppo, Rick Ramirez, Mark Koester, Siok Leong, Zachary F. Mattes, Erin Gallagher, Binh Lee, Franco Abbate, Lila Ghamsari, Gene Merutka, Abi Vainstein-Haras, Barry J. Kappel, Jim A. Rotolo

## Abstract

Glioblastoma (GBM) is the most prevalent primary brain cancer, with poor prognosis and limited therapeutic options available. The genetic and cellular heterogeneity characteristic of GBM contributes to poor response rates. Activating mutations of the epidermal growth factor receptor (EGFR) gene are among the most frequent alterations in GBM, occurring in roughly half of cases. Despite the prevalence of EGFR mutations, EGFR inhibition has shown limited success in GBM. The transcription factor C/EBPβ is a master regulator of the mesenchymal transformation in GBM, an aggressive state characterized by increased invasiveness and resistance to chemotherapy. Lucicebtide is a C/EBPβ antagonist peptide with demonstrated single agent activity in patients with recurrent GBM that is currently being evaluated in a clinical trial in combination with radiation and temozolomide in patients with newly-diagnosed GBM (NCT04478279), with emerging data supporting clinical activity in that setting. Here we show that in the TCGA-GBM dataset, patients with EGFR mutations display significant enrichment of a high C/EBPβ activity signature. Functionally, genetic inactivation of EGFR by CRISPR results in synthetic lethality in the presence of lucicebtide in GBM cell lines, and synergistic in vitro cytotoxicity and suppression of C/EBPβ target gene expression was observed in combination experiments with lucicebtide and EGFR inhibitors. Finally, enhanced anti-tumor activity was demonstrated in vivo in the combination setting, as combined subpharmacologic dose levels of lucicebtide and the EGFR inhibitor osimertinib potently suppressed GBM xenograft growth. These data identify EGFR and C/EBPβ dependencies in GBM and support lucicebtide combination with EGFR inhibitors as a potential therapeutic option for a sizable fraction of GBM patients.

## Introduction

Glioblastoma (GBM) is the most common and aggressive malignant brain tumor, with a 5-year survival of 6.8% (1) and median overall survival of 14.6-17 months (2). Standard of care therapy includes maximal surgical resection, followed by postoperative radiation therapy along with the alkylating agent temozolomide (TMZ) (3). Frequent genetic lesions found in GBM include activating mutations of Epidermal Growth Factor Receptor (*EGFR*) and Phosphoinositide 3-Kinase (*Pi3K*), mutations in Telomerase Reverse Transcriptase (*TERT*) promoter and inactivating mutations in the tumor suppressors *TP53*, *PTEN* and *CDKN2A/B* (4). *EGFR* genetic alterations include multiple non-exclusive mechanisms, including copy number amplifications, point mutations, fusions, and in-frame deletions which include the recurrent *EGFRvIII,* an in-frame deletion lacking exons 2-7 (4,5). In GBM, it is estimated that ∼50% of cases carry EGFR alterations resulting in an aberrantly active receptor. Consequently, EGFR was heavily pursued as a target in the clinic by a variety of approaches, including small molecule tyrosine kinase inhibitors (TKIs) (6), monoclonal antibodies (7), antibody drug conjugates (ADCs) (8) and chimeric antigen receptor T cells (CAR-T) (9). To date, EGFR inhibition has shown limited success in GBM, likely due a combination of factors including cellular heterogeneity of the disease, poor blood brain barrier (BBB) penetration for some compounds, intrinsic resistance arising from compensatory signaling pathways, and acquired resistance due to secondary EGFR mutations (6,10,11).

CCAAT/Enhancer Binding Protein β (C/EBPβ) is a developmental stage transcription factor of the C/EBP family with functions in cell proliferation, control of cell fate in several tissues, and innate immune regulation (12). In addition, C/EBPβ was identified as a master regulator of the mesenchymal transformation in GBM (13), a tumor cell lineage associated with aggressive disease and therapeutic resistance. A bidirectional loop between EGFR and CEBPB in GBM has been described where each factor transcriptionally regulates the other (14). Lucicebtide (formerly known as ST101) is a C/EBPβ antagonist peptide that resulted in durable responses in newly diagnosed and recurrent GBM patients in phase 2 clinical trials (15). Lucicebtide is known to act through distinct mechanisms; direct suppression of cancer cell growth (16), disruption of mesenchymal transformation programs in GBM, and activation of anti-tumor immune responses by reprogramming immunosuppressive macrophages in the tumor microenvironment (TIME) toward an immune-active program (17,18). The impact of lucicebtide on EGFR signaling has not been previously investigated.

Here we sought to uncover and validate combination strategies with lucicebtide for genetically identifiable GBM cases. Our strategy consisted of leveraging the publicly available TCGA-GBM cohort that has been extensively characterized by transcriptional and genetic profiling (4). We then aimed to validate synergistic interactions with lucicebtide by genetic and pharmacologic approaches aimed at inhibiting EGFR.

## Material and Methods

### Peptide Synthesis

Lucicebtide (*H_2_N*-vaeareelerlearlgqargelkkwkmrrnqfwlklqr-*OH*, lot number 30-19-0226-01) was synthesized by Fmoc solid-phase peptide synthesis (SPPS) and the mass and sequence were confirmed by mass spectrometry. Lucicebtide solution was prepared from lyophilized powder in sterile milli-Q H_2_O containing 270 mM trehalose to a stock concentration of 2 mM.

### Lucicebtide activity signature derivation

RNAseq profile of U251 cells treated with lucicebtide for 24 hrs (GSE213013) (16) were processed by removing genes with less than 2 FPKM in 2 or less samples. Expression levels were log2-transformed and unsupervised clustering (UC) was performed using genes with standard deviation greater than 2 (MatCalc (19)). After removing Y RNA, the list of genes downregulated by lucicebtide were pared down to a 37 genes signature that comprises the lucicebtide activity signature (LAS; list of genes in Supplementary Table 1).

### TCGA cohort classification

RNAseq and mutational profiles of the TCGA-GBM dataset (n=293) were downloaded from the GDC portal. For samples with duplicate RNAseq profiles, the first sample was used for analysis. Samples were then stratified into the high LAS set if they scored in the top quartile (n=74) or low LAS set for the bottom three quartiles, n=219). Two-sided Fisher T-test for the mutational status was performed using the Python scipy package. Supplementary Table 5 from Brenner et al. (4), was used as source of annotation of specific EGFR alterations (amplification, mutation, structural changes). The LAS classification was generated after removing cases for which EGFR status information was not available. For copy number data, copy gains and euploid state were both considered in the “Not Amplified” category. Supplementary Table 2 reports mutational status for genes listed in Figure 1C LAS score for the TCGA-GBM cases.

**Figure 1.**
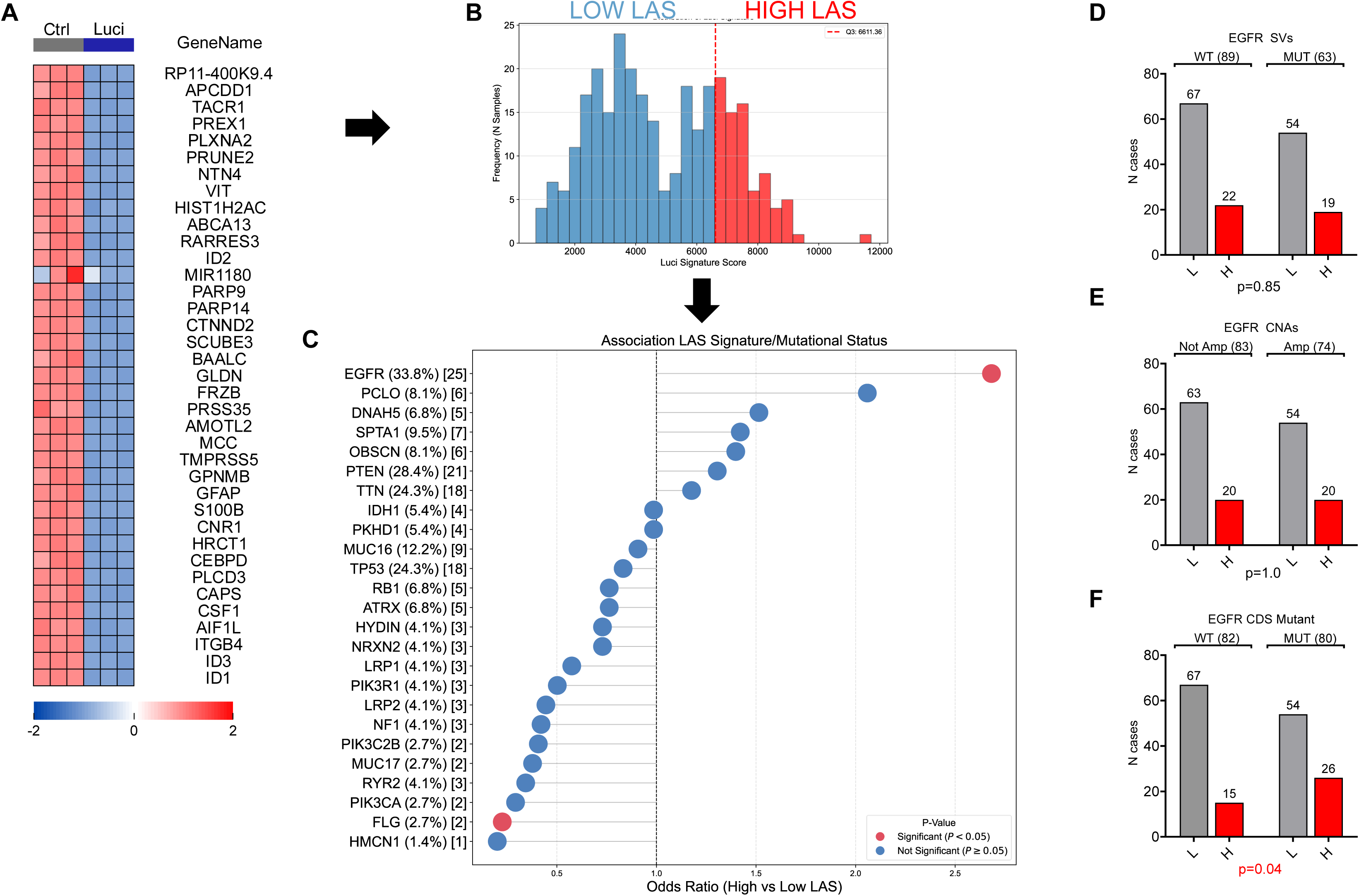
EGFR mutations are associated with lucicebtide activity signature. **A)** Heat map for expression level of the 37 genes included in the lucicebtide activity signature (LAS) in U251 cell lines untreated (Ctrl, control) or treated with lucicebtide for 24hrs. **B**) Histogram for TCGA-GBM cases (n=293) classified into low or high LAS using a Q3 threshold (red dotted line). **C**) Lollipop chart for Fisher contingency test for 25 frequently mutated genes in GBM (defined as mutated in >5% cases). X-axis indicates odd-ratio for high vs low LAS. Color-coding indicates significance level. (Red, p<0.05, Blue, p>0.05, Fisher T-test). Y-axis reports gene name, frequency of mutations and number of mutated cases in the high LAS subset. **D-F**) Contingency analysis for EGFR structural variants (SV, **D**, n=162), copy number alterations (CNAs, **E**, n=157) and coding-sequence changing mutants (CDS, **F**, n=162) with LAS status (grey, low LAS; red; high LAS). Number of cases for each category is indicated above the corresponding bar. P-values represent Fisher T-tests.

### Cell culture

T98G cells (ATCC, CRL-1690) were maintained in complete EMEM 10% (ATCC 30-2003), and U251 cells (Sigma, 9063001) were plated in complete MEM 10% (Thermo Fisher 11095098). Lenti X cells (Takara Bio USA, 632180) were cultured in complete DMEM 10% (Thermo Fisher, 11965092).

### Lentiviral transduction and CRISPR

CAS-9 expressing lentivirus was packaged in Lenti X according to the manufacturer’s instructions. Briefly, Lenti X cells were transfected with CAS9 plasmid and packaging vectors (Applied Biologics Materials) at a ratio of 5:1:1:1. Supernatant was collected at 48 hours and incubated O/N at 4°C with Lenti-X Concentrator (Takara) at a 1:3 ratio. Lentiviral concentrate was resuspended in 300 µL, aliquoted and stored at −80°C. To establish CAS9 stably expressing cells, T98G and U251 cells were plated in 6-well plates on Day 0 at 2E5 cells/well with 2 mL media/well. For each cell line and infection, 3 wells were seeded. On Day 1, following 16 hours seeding, 30uL of CAS9 virus concentrate was added to each well and supplemented with 10 uL Polybrene (final concentration (4 ug/mL, Santa Cruz Biotechnology). Six-well plates were then spinoculated at RT for 45 minutes at 2,000 rpm. On Day 4, cells were trypsinized and seeded into a T25 in 5 mL of the corresponding media supplemented with 10 uM Blasticidin HCl (Goldbio). On Day 11, cells were trypsinized and assessed for CAS9 expression by Western Blot. CRISPR sgRNAs were chosen from the VBC database prediction as the top ranked target sequences targeting different exons (20). Two independent target sequences for each target gene were used (EGFR_A: GATGTTCAATAACTGTGAGGTGG; EGFR_B: GATGTGGAGATCGCCACTGATGG; and PCNA: ATACGTGCAAATTCACCAGAAGG) and cloned in sgRNA expressing vectors (Cellecta, SVCRU6EGP-L, SVCRU616-L). For control, a neutral sequence previously identified was used (21). Infection and selection for the sgRNA vectors followed the previously described protocol. Selection was completed through Puromycin (Sigma) at 2.5 µM in T25 cultures for 3 days. Target suppression was confirmed by Western blot.

### CRISPR synthetic lethal competition screen

GFP-tagged, puromycin selected T98G and U251 knockout lines for PCNA, EGFR or control were mixed at a 1:1 ratio with TagRed-tagged CRISPR control cells in 12-well plates at 2.5E4 cells/mL each for 2 mL total volume (5E4 total cells/well). Cell mixtures were acquired on Day 0 on a MACSQuant flow cytometer (Miltenyi), to verify equal representation of the two cell populations. On day 1, media was aspirated and replaced with fresh complete media containing 0, 2.5, 5, or 10 µM lucicebtide. On day 7, cells were tryspinized, transferred to a 96-well deep well and the eGFP:TagRed ratio was determined by flow cytometry using a MACSQuant. Flow cytometry data were analyzed using FlowJo and statistics were generated using GraphPad PRISM. EGFP gates were manually adjusted for each sgRNA population to accommodate for variability in eGFP expression among different sgRNAs constructs.

### Western Blot

Cell lines were pelleted and lysed using RIPA buffer (Thermo Fisher). The cell lysates were quantified using Pierce BCA Protein Assay Kit (Thermo Fisher) according to the manufacturer’s instructions. Samples were mixed with NuPage LDS Sample Buffer and NuPage Sample Reducing Agent (Thermo Fisher), boiled for 5 minutes at 95°C, loaded into a NUPAGE 4-12% Bis-Tris Gel or Novex 4 to 12% Tris-Glycine Gel (Thermo Fisher), and subjected to SDS-PAGE. The samples were transferred to a PVDF membrane filter (Bio-Rad) using a semi dry system transfer (Bio-Rad) or a XCell II Blot Module wet transfer system (Thermo Fisher). Blots were then incubated in 5% milk (Bio-Rad) or BSA for 1 hour at RT for blocking. Primary antibodies used were, Phospho-C/EBPB (Thr235) (1:1000, Cell Signaling Technology #3084), C/EBPB (E2K1U) (1:1000, Cell Signaling Technology #43095) and Anti-Beta-Actin-Peroxidase antibody, (1:10000, Sigma, #A3854). The secondary antibodies used were anti-mouse IgG, HRP-linked (1:2000, Cell Signaling) and anti-rabbit IgG, HRP-linked Antibody (1:2000, Cell Signaling). HRP-actin signal was developed directly. Blots were developed using SuperSignal West Pico PLUS Chemiluminescent Substrate (Thermo Fisher). For blot reprobing, blots were stripped using Restore PLUS Western Blot Stripping Buffer (Thermo Fisher), re-blocked and incubated with primary antibody as indicated.

### Viability assays

T98G and U251 cells were plated at 2.5E3 cells/well on Day 0 in 24 96-well plates (Corning). On Day 1, EGFR inhibitors (osimertinib, afatinib or BTDX1535 (Selleck)) and lucicebtide were added in serial dilution to the plate according to the indicated concentrations. DMSO concentration was normalized across the whole plate. Each assay was performed in eight replicates per combination. Of the eight replicates, four were read out using TiterGlo (Promega) via a SpectraMax plate reader according to the manufacturer’s instructions, and four were subject to an Annexin-V-based flow cytometry assay for cell viability. For Annexin-V staining, cells were washed in PBS, trypsinized and transferred to a 96-well deep plate (USA Scientific). Cells were washed twice at 4°C with BioLegend Cell Staining Buffer (BioLegend) and resuspended in Annexin-V Binding Buffer (BioLegend). For every 1E6 cells, 5 µL of Annexin-V pacific blue (Biolegend, 640918) and 1 µL of Sytox Deep Red Nucleic Acid Stain (BioLegend, S11380) were added. After gentle vortexing, cells were stained for 15 minutes at RT protected from light, washed and acquired on a MACSquant 16 instrument. Analysis was performed using the Combenefit software package (22). Dose response curves were fitted using GraphPad software (Dotmatics) by the Find ECanything function and least square regression method. Areas Under the Curve (AUCs) were calculated using a 4-PL logistic from the package scipy. Errors were estimated by bootstrap (1000 iterations).

### Quantitative RT-PCR (targets)

Cells were treated with lucicebtide and osmeritinib combinations as previously described, before being pelleted and lysed with Trisol for RNA extraction following manufacture’s protocol. The RNA quality and quantity for each sample were measured by Nanodrop. The cDNA was then generated from the RNA samples through the SuperScript IV VILO Master Mix with ezDNAse enzyme (Thermo Fisher) following manufacturer’s protocol. The QuantStudio 6 Flex real-time thermal cycler and Fast SYBR Green Master Mix (Thermo Fisher) were used for qPCR. Samples were run using 100 ng cDNA in quadruplicates in 384-well plates with primers listed in Supplementary Table 4. Control reactions without reverse transcriptase were performed. Log2 normalized expression (2^ΔΔCt) and standard error of mean were used to determine fold change of expression between treated and untreated samples in relation to β-actin housekeeping gene (23). Statistical significance between groups was determined using Student’s T-test.

### U251 xenotransplant model

Five million U251 cells were mixed 1:1 with Matrigel (Corning) and injected in the right flank of 5-6 week old Nu/J female mice. One week after injection, mice with detectable tumors were randomly assigned to treatment or control cohorts so that mean of tumor volume was within one SEM for all cohorts. Tumors were tracked by bi-weekly caliper measurements and volumes calculated as previously described (17). For the combination experiment, lucicebtide (10mg/kg) was administered three times weekly; osimertinib (2mg/kg) was administered s.c. daily. Controls were administered 10% DMSO in PBS. All aspects of animal care were in accordance with the Guide for Care and Use of Laboratory Animals and all experiments were approved by the Institutional IACUC at New York Medical College (NYMC).

## Results

### A GBM lucicebtide activity signature is associated with EGFR genetic activation

A lucicebtide activity signature (LAS) was defined using RNAseq profiling of the U251 GBM cell line treated with lucicebtide for 24hrs as described (16). Unsupervised clustering identified a set of 37 genes downregulated by lucicebtide (**Fig 1A**, **Supplementary Table 1**), including previously known lucicebtide targets such as ID1, ID3 and GFAP (16). The LAS signature was then used to classify the 293 TCGA-GBM cases with available RNAseq profiles into ‘Low’ and ‘High’ LAS sets using a Q3 threshold (17) (**Fig. 1B).** The association of mutational status of the 25 most frequently mutated genes in GBM patients (mutated in more than 5% of the TCGA-GBM dataset) with high LAS signature was determined for each gene by Fisher contingency test between mutational and LAS status. EGFR mutations were the only alterations found to be significantly overrepresented in high LAS cases (**Fig. 1C**, **Supplementary Tables 2-3**). Specifically, in the low LAS set, 16% of GBM cases (35 out of 219) carried EGFR mutations, while in the high LAS set, the frequency of cases increased to 34% (25 out of 74; P<0.002, 2.7 odds-ratio). Cases carrying mutations in the FLG gene were significantly enriched in the low LAS subtype (11% vs 2.7% in the high LAS set), although its role in GBM pathogenesis is unclear.

EGFR may be genetically altered multiple ways in GBM, including structural variants (SV) encoding in-frame deletions and translocations, copy number alterations (CNA), and point mutations within the coding DNA sequence (CDS) (4). To understand if high LAS is linked to a specific category of EGFR alterations, we performed contingency tests between EGFR status as annotated in the Brennan study (4) and LAS classification for the subset of samples for whom genetic information was available (SV and CDS, n=162; CNA, n=157). Notably, EGFR CDS mutations were significantly associated with high LAS, while SV and CNA samples were not (**Fig. 1D-F**). Overall, these data identify that LAS is distinctively associated with EGFR mutations, and suggests that EGFR-dependency may be linked to C/EBPβ activity in GBM.

### Genetic EGFR suppression is synthetically lethal with lucicebtide treatment in GBM cells

To functionally test if EGFR and C/EBPB dependencies are related, we generated isogenic CRISPR knockout pools in the T98G and U251 GBM lines and tested them in a competition assay in the presence of lucicebtide or control (**Figure 2A**). To do this, EGFP-tagged pools targeting a neutral genome region (Ctrl) or EGFR (two pools with distinct sgRNAs, denoted EGFR A and EGFR B) were mixed at 1:1 ratio with a tagRed-infected Ctrl pool and treated with increasing concentrations of lucicebtide for one week. EGFR sgRNA produced nearly complete suppression of EGFR protein expression as assessed by Western Blot in both cell lines (**Figure 2B,D**). The normalized EGFP:tagRed ratio was then used to measure the fitness of EGFR knockout cells exposed to lucicebtide compared to control cells under the same conditions. The fitness of EGFR knockout cells substantially decreased in the presence of lucicebtide, reaching ∼70% and ∼60% depletion (****p<0.0001) compared to untreated controls in T98G EGFR A and EGFR B respectively (**Figure 2C**; Statistics, 2-way ANOVA T-test with Dunnett’s correction). Similar levels of synthethic lethality were observed in U251 cells (**Figure 2E**). In control cells, lucicebtide exposure decreased cell viability in a dose-dependent manner consistent with published results (16), however no differences in the control mixture ratio were observed in either T98G or U251 cell lines. These data indicate that sensitivity to lucicebtide is enhanced by EGFR depletion and point to a targetable co-dependency of the EGFR/C/EBPβ axis.

**Figure 2.**
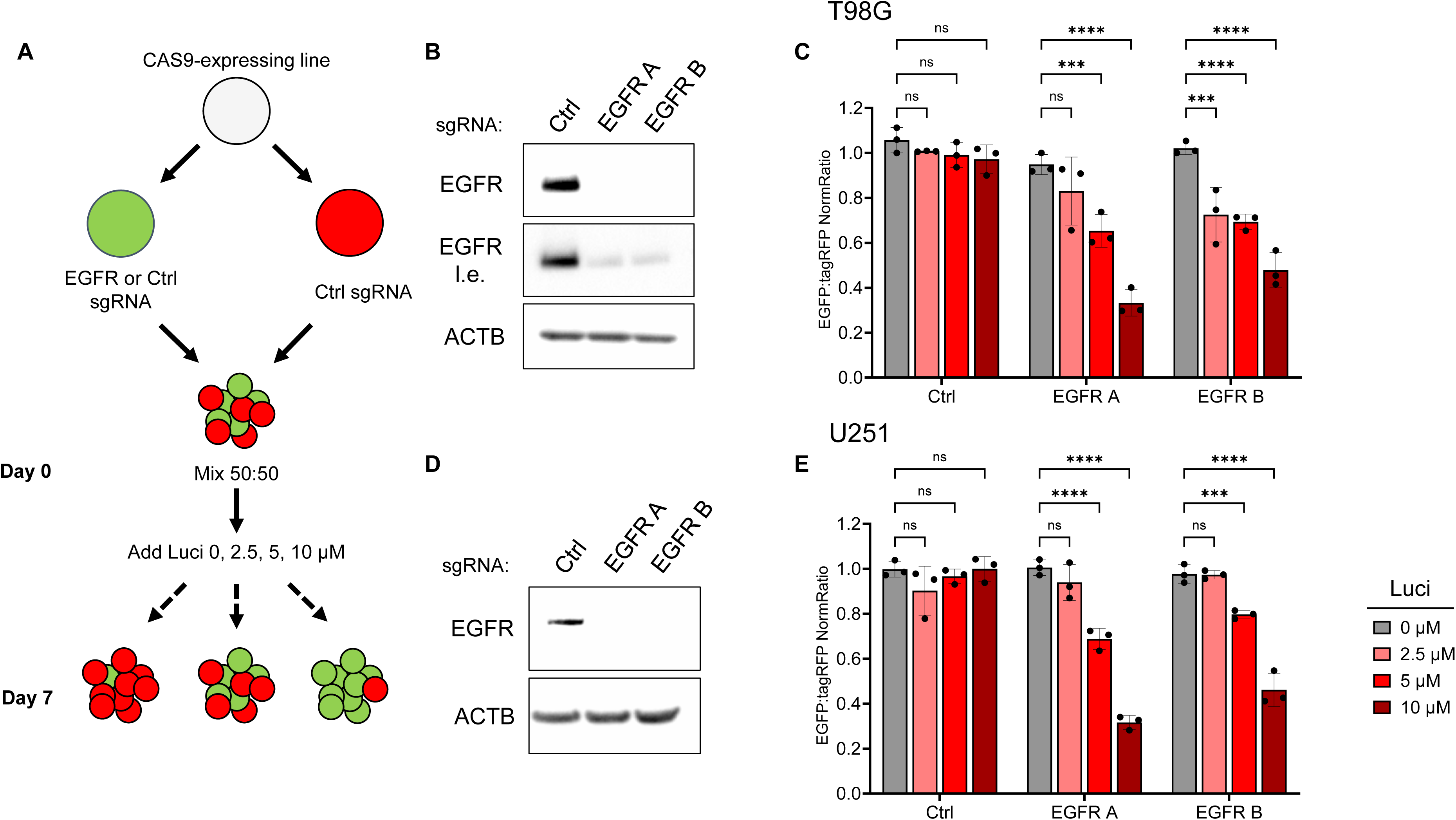
Synthetic lethality of EGFR suppression with lucicebtide in GBM. **A)** Outline of the CRISPR synthetic lethal approach. CAS9-expressing GBM lines were transduced with EGFP-tagged sgRNAs targeting a control region (Ctrl) or two non-overlapping EGFR sgRNAs (EGFR A and B). Cells were mixed with TagRed-expressing Ctrl sgRNA cells and plated in the presence of increasing concentrations of lucicebtide. After one week, cell mixtures were recovered and assessed for EGFP:TagRed cell ratio. **B,D)** Western blot assessing suppression of EGFR in selected T98G (**B**) and U251 (**E**) control, EGFR A and EGFR B pools. β-Actin was used as loading control. l.e., long exposure. **C, E)** EGFP:TagRed Normalized Ratio for T98G (**C**) or U251 (**E**) cells infected with control (Ctrl), EGFR A or EGFR B sgRNAs and treated with 0, 2.5, 5 or 10 µM lucicebtide. Assays were performed in triplicate. Error bars represent standard deviations. Statistics: 2-way ANOVA, T-test with Dunnet’s correction for the indicate comparisons (*** p<0.0001; *** p<0.001; ns, not significant).

### ST101 synergizes with EGFR inhibitors

To test if pharmacological EGFR inhibition impacts lucicebtide sensitivity in GBM cells, we performed checkerboard assays of lucicebtide with afatinib, osimertinib, and BDTX-1535, three distinct advanced-generation EGFR TKIs (24–26). For these assays, T98G and U251 cells were treated with lucicebtide concentrations ranging from 0 to 2.5 µM, each of the EGFR TKIs ranging from 0 to 10 µM, or the combination of lucicebtide and TKI. Dose responses were measured at 48hrs by Annexin V/Sytox Red flow cytometry assay and in parallel by CellTiter-Glo, and synergy was measured using Combenefit by the BLISS method (22,27). Lucicebtide was potently synergistic in combination with all three inhibitors in both assays (**Figure 3A-F**). To quantitate the effect of the synergy, dose response curves were fitted and the osimertinib EC_10_ (T98G) and EC_20_ (U251) were evaluated with and without lucicebtide. These effective concentrations were chosen because the maximum effect achieved by 10 µM osimertinib alone did not reach EC_50_. (**Figure 3 G-L**). Data indicates that 1.25 µM lucicebtide induced a 1.9-fold decrease in osimertinib EC_20_ in T98G and a 3.2-fold decrease in EC_10_ in U251 (**Figure 3H,K)** compared to osimertinib alone. Accordingly, 1.25 µM lucicebtide resulted in a 35% decrease in the area under the curve (AUC) in T98G cells and a 25% decrease in the U251 line (**Figure 3I,L**). Consistent results were observed using CellTiter-Glo readouts (not shown). These results demonstrate that pharmacological EGFR inhibition is synergistic with lucicebtide and suggests the feasibility of lucicebtide-EGFR TKI combination strategies for GBM patients with EGFR mutations.

**Figure 3.**
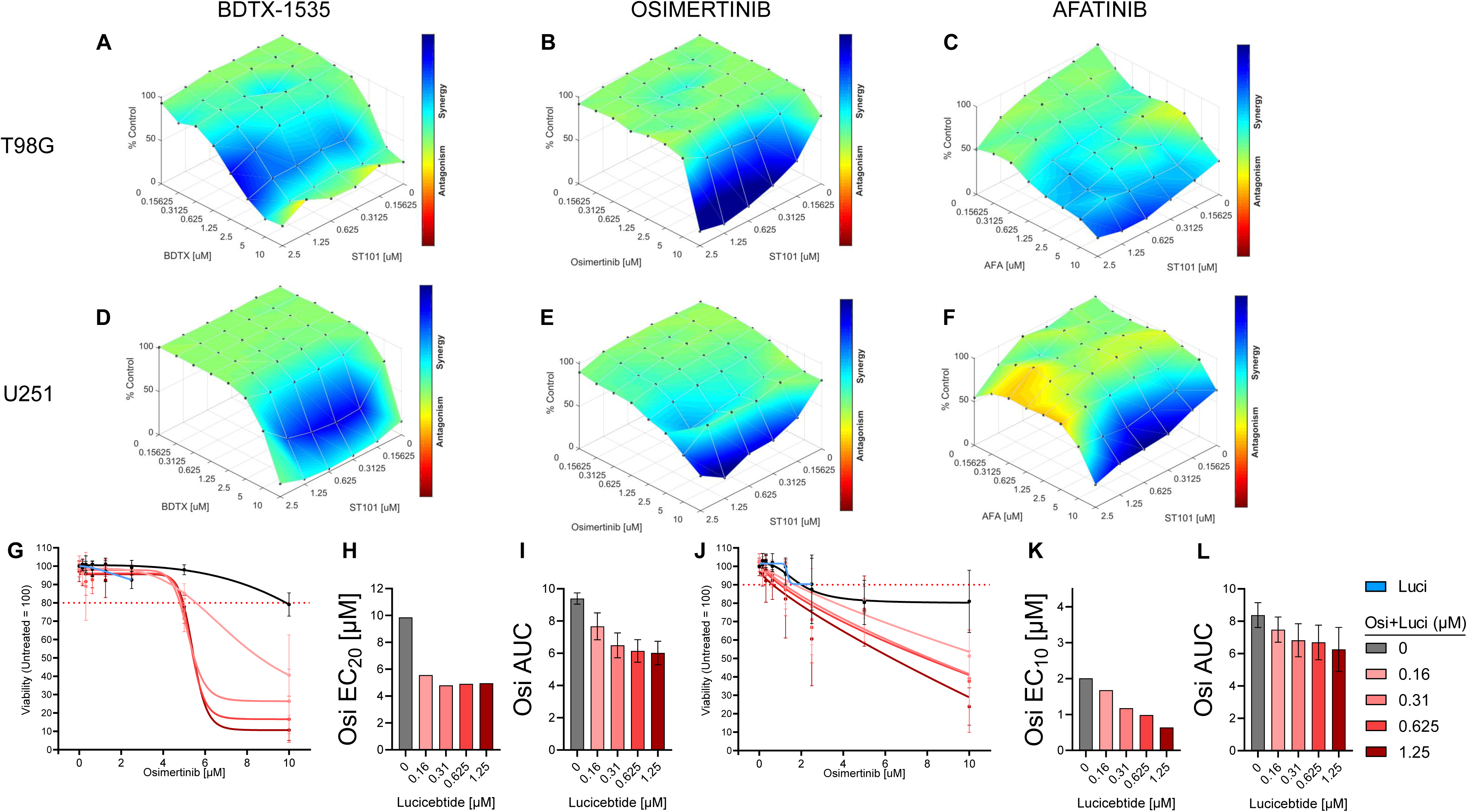
EGFR inhibitors and lucicebtide synergize in GBM cell lines. **A-F)**Bliss surface models for two GBM lines (T98G, U251) treated with lucicebtide and three chemically distinct EGFR inhibitors (BDTX-1535 (**A,D:** BDTX), osimertinib (**B,E;** OSI), afatinib (**C,F:** AFA)) at the indicated concentrations and combinations. Viability was assayed at 48hrs by an Annexin V and Sytox Red apoptosis-based assay (see Methods). All points were assayed in 4 independent replicates. **G-L**) Dose response curves, absolute effective concentrations (ECs) and AUCs bar graphs for osimertinib combined with the indicated lucicebtide concentrations in T98G (**G-I**) and U251 (**J-L**) lines. For each cell line an appropriate EC was utilized depending on the max effect reached in the osimertinib-only response (T98G, EC_20_ (**H**); U251, EC_10_ (**J**)). For dose-response graphs, dotted lines represent EC_20_ (**G**,T98G) or EC_10_ (**I**, U251) levels. AUCs (T98G, **I**; U251, **L**) were integrated over the 0-10 µM linear osimertinib range. Error bars for AUCs represent standard deviations obtained by bootstrapping (1000 replicates).

### C/EBPβ is targeted by osimertinib and lucicebtide via non-overlapping mechanisms

Previous studies identified that disruption of C/EBPβ dimerization with lucicebtide resulted in its degradation in a proteosome-dependent manner (16). Consistent with previous findings, exposure of T98G cells to 10 µM lucicebtide was sufficient to enhance C/EBPβ degradation. While osimertinib did not result in significant impact on total C/EBPβ protein levels, a substantial reduction was observed in C/EBPβ Threonin-235 phosphorylation, a MAPK-dependent site that has been demonstrated to be critical for *CEBPB* promoter transactivation (28) (**Figure 4A**). Thus, it was hypothesized that the synergy observed with the lucicebtide and osimertinib combination may be due to parallel impact on C/EBPβ. Supporting this hypothesis, *ID1* transcript levels as a read-out for C/EBPβ activity were unchanged in subpharmacologic lucicebtide- or osimertinib-only treated cells, but significantly suppressed in combination treated cells compared to untreated or single-treatment conditions (**Figure 4B**). These data indicate that the combination activity of lucicebtide and osimertinib converge, at least in part, on C/EBPβ transcriptional activity.

**Figure 4.**
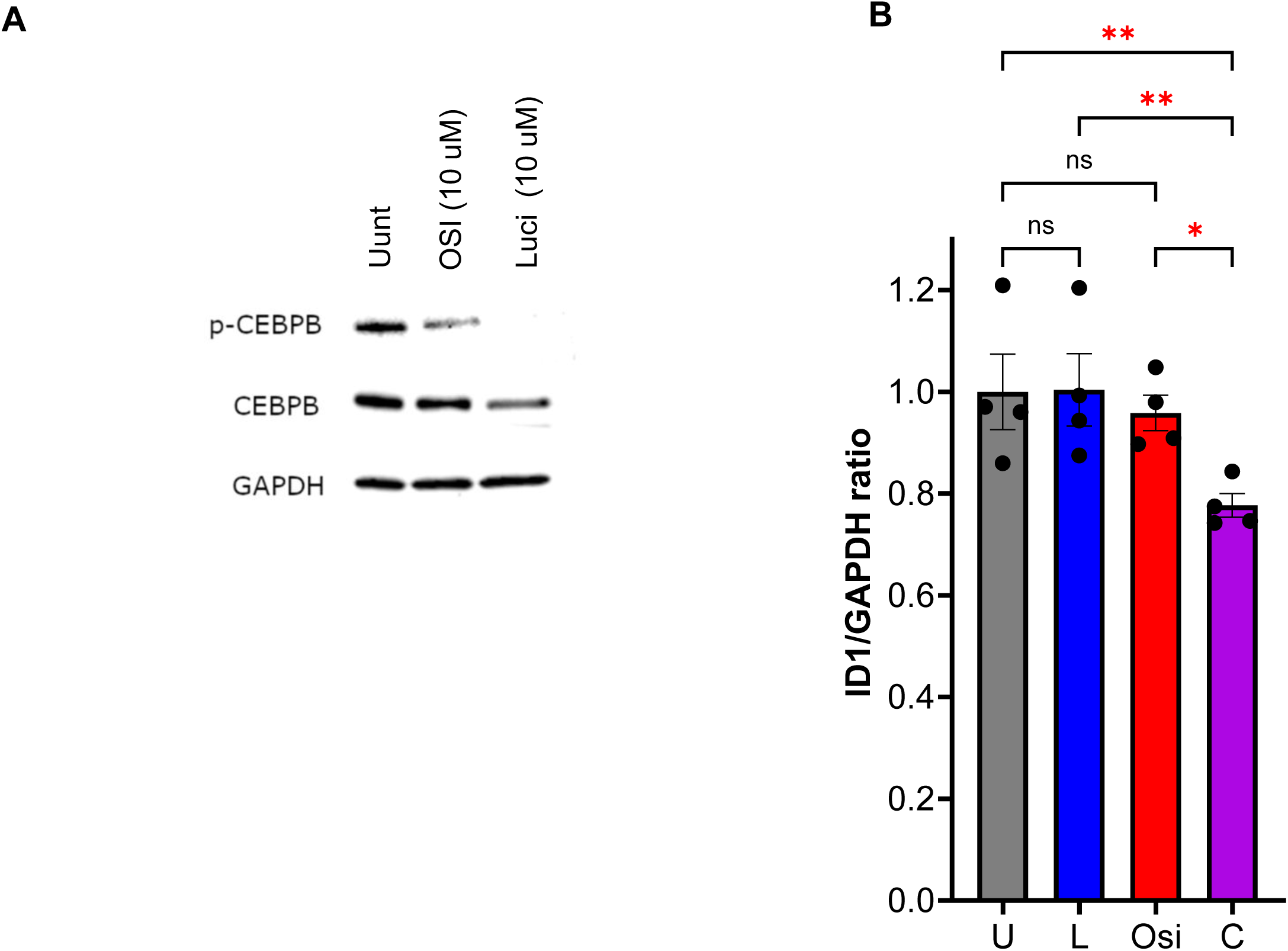
EGFR inhibition and lucicebtide converge on suppression of C/EBPβ transcriptional activity. **A**) Western Blot for total and phospho-C/EBPβ in the indicated conditions. **B**) QPCR analysis for ID1/Actin transcript in the indicate conditions (U, untreated; L, lucicebtide 0.625 uM; O, osimertinib 5 uM, C, combination). Statistics, 1-way ANOVA (**p<0.01, ns, not significant).

### Combination of osimertinib and lucicebtide results in enhanced anti-tumor activity in U251 xenotransplants in vivo

To investigate combination efficacy of lucicebtide and osimertinib in vivo in the U251 GBM xenograft model, tumor-bearing nude mice were treated with a subpharmacologic dosing regimen for osimertinib (osimertinib dose-response in U251 model shown in **Supplementary Figure 1**) and subpharmacologic lucicebtide (16). Specifically, mice were treated with vehicle as control (10% DMSO in PBS), subpharmacologic osimertinib (2.5 mg/kg QD), subpharmacologic lucicebtide (10 mg/kg 3x/week), or subpharmacologic lucicebtide in combination with osimertinib, and tumor volumes were evaluated every 3 days for 30 days. Subpharmacologic dosing of either compound alone did not yield a significant difference in tumor volumes vs. control (**Figure 5A**, control vs osimertinib, p=0.71; control vs lucicebtide, p=0.54). Conversely, the lucicebtide and osimertinib combination resulted in 73% tumor growth inhibition (TGI) at day 30 compared to control (p=0.0007), and 66% and 68% TGI compared to monotherapy (combination vs. osimertinib, p=0.006; and combination vs. lucicebtide, p=0.011). None of these treatment regimens significantly impacted relative body weight during the course of the treatment (**Supplementary Figure 2**). Combination efficacy was attributed to direct activity on tumor cells, as minimal host macrophages are present in this model (data not shown). These in vivo data are consistent with in vitro genetic and pharmacologic checkerboard assays, and support the combination of lucicebtide and EGFR inhibition as a potential therapeutic option for the population of GBM patients that harbor EGFR mutations.

**Figure 5.**
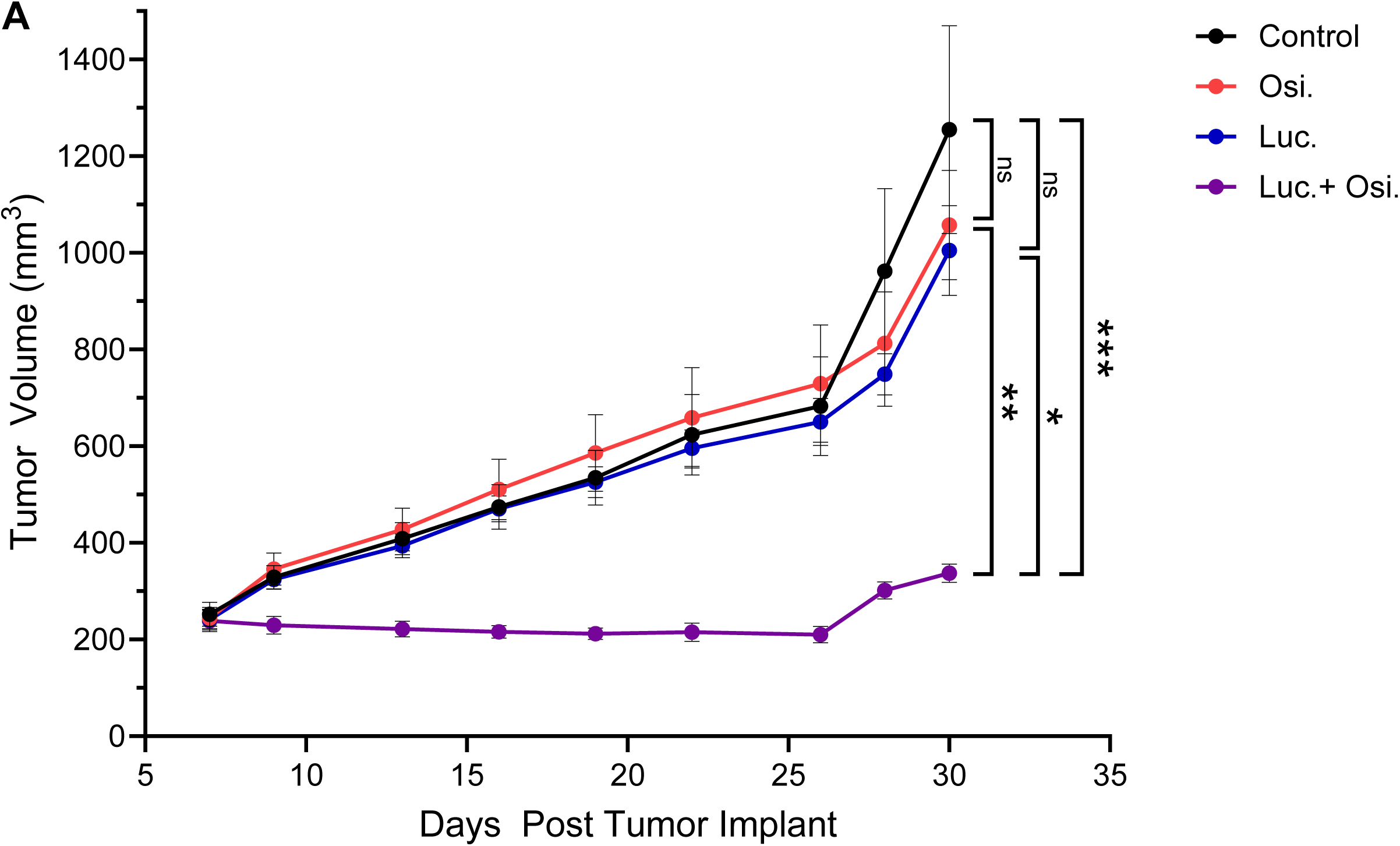
Combination of lucicebtide and osimertinib enhances anti-tumor activity in GBM in vivo. **A)** Tumor volumes of mice transplanted with U251 cells and treated with Control (vehicle, black) or with the indicated treatment (subpharmacologic osimertinib, 2 mg/kg QD, red; subpharmacologic lucicebtide 10 mg/kg 3x/week, blue; combination, purple). Statistics indicate 1-way ANOVA (***,p<0.001;**,P<0.01; *,p<0.05; ns, not significant; n=5/group) for volumes at Day 30. Error bars represent standard deviations.

## Discussion

GBM is a hard-to-treat cancer with poor life expectancy and few clinical options. Genomic profiling efforts have identified that EGFR alterations, in the form of activating in-frame deletions (*EGFRvIII*), amplifications or sequence-changing mutations, are one of the most frequent genetic events in GBM patients, with an estimated 50% occurrence in the TCGA-GBM cohort. Despite the prevalence of genetic alterations, EGFR inhibition as a single-agent strategy has failed to achieve clinical impact, likely for the complex dependencies and cellular heterogeneity of these cancers. Here we discovered an association in GBM between EGFR mutations and the expression of a transcriptional signature impacted by lucicebtide. This association was specific, as none of the other 25 most frequently altered genes in GBM correlate with the lucicebtide active signature.

Our data identified a functional co-dependency between EGFR and C/EBPβ in GBM, such that EGFR deletion by CRISPR or inactivation via three distinct EGFR TKIs is synergistic with lucicebtide. EGFR inhibition led to a reduction in the active phospho-T235 C/EBPβ protein, a mechanism that is distinct but potentially complementary to lucicebtide antagonism of C/EBPβ dimerization, which prevents C/EBPβ activity and targets the protein for degradation (16). These data suggest that the combination efficacy observed occurs via distinct mechanisms that converge upon modulating C/EBPβ activity. Consequently, quantitative RT-PCR showed that the lucicebtide-EGFRi combination at concentrations that are not effective as single-agents significantly suppressed expression of *ID1*, a previously described C/EBPβ target gene and prominent factor involved in epithelial to mesenchymal transformation (29).

Clinically, GBM tumors are supported by a rich TME characterized by a high percentage of immune-suppressive macrophages (30). We have previously shown that lucicebtide reprograms macrophages toward immunoactivity in immune checkpoint inhibitor-resistant tumors, enhancing the anti-tumor activity of anti-PD1 in these models. Literature suggests that *EGFRvIII* mutations promote immunosuppressive macrophage polarization and support GBM growth, however the impact of EGFR inhibitors on macrophage polarization is not described (31). While potent in vitro synergy was observed with the lucicebtide and EGFRi combination in GBM xenografts, notably these models do not recapitulate the contribution of the TME to tumor growth. Thus, in models were the TME is actively involved in tumor progression, additional synergies between lucicebtide and EGFR inhibitors may be identified.

Overall, our data suggests that combination with lucicebtide may be a path to improve the overall clinical benefit of EGFR inhibitors in GBM. Second and third-generation EGFR TKIs have raised particular interest in GBM, as inhibitors such as afatinib (25) and osimertinib (26) have clinical activity on brain metastases in non-GBM EGFR-driven malignancies (32) and BDTX-1535 has shown the ability to penetrate the BBB (24). Additionally, latest-generation TKIs have been designed to overcome *EGFR* gate-keeper mutations that frequently drive acquired resistance (33). Inferring that *EGFR* mutations impact GBM by enhancing C/EBPβ activity, as supported by the high LAS in these patients, the lack of single-agent EGFR inhibitor activity may be due, at least in part, to EGFR-independent C/EBPβ activity. As several TKIs have already been approved for other oncology indications, combination with lucicebtide, a peptide demonstrated in the clinic as safe and active in GBM patients, presents an opportunity to effectively use EGFR inhibitors in GBM and enhance patient benefit. Finally, as EGFR alterations represent an established, high-frequency GBM genetic hallmark, this study provides a putative biomarker strategy to identify GBM patient subsets who may benefit from combination therapies with lucicebtide.

## Acknowledgements

The authors wish to thank all the Sapience Therapeutics team for supporting this work.

## Contributions

J.D., C.S. and J.R. designed the study and wrote the manuscript. J.D., C.S., and R.R. performed experiments, developed reagents and analyzed data. S.L., M.K.. E.G., B.L., F.A. and Z.F.M performed experiments. B.K. and A.V.H. contributed to the study design. J.R. supervised the study. All the authors reviewed and edited the manuscript.

## Data Availability Statement

The GBM transcriptional profiles, mutational and survival data analyzed in this study were obtained from the TCGA repository (portal.gdc.cancer.gov/projects/TCGA-GBM). The lucicebtide GBM signature is listed in Supplementary Table 1 and derived from GSE213013. Other data generated in this study are available within the article or are available upon reasonable request from the corresponding author.

## Supplementary Materials

**Supplementary Table 1.** List of genes included in lucicebtide activity signature (**LAS**).

**Supplementary Table 2**. Contingency values and Fisher T-test statistics for each frequently mutated GBM gene and LAS status.

**Supplementary Table 3**. Mutational annotation and LAS score for the TCGA-GBM cohort.

**Supplementary Table 4**. Primer list used in this study.

**Supplementary Figure 1.**
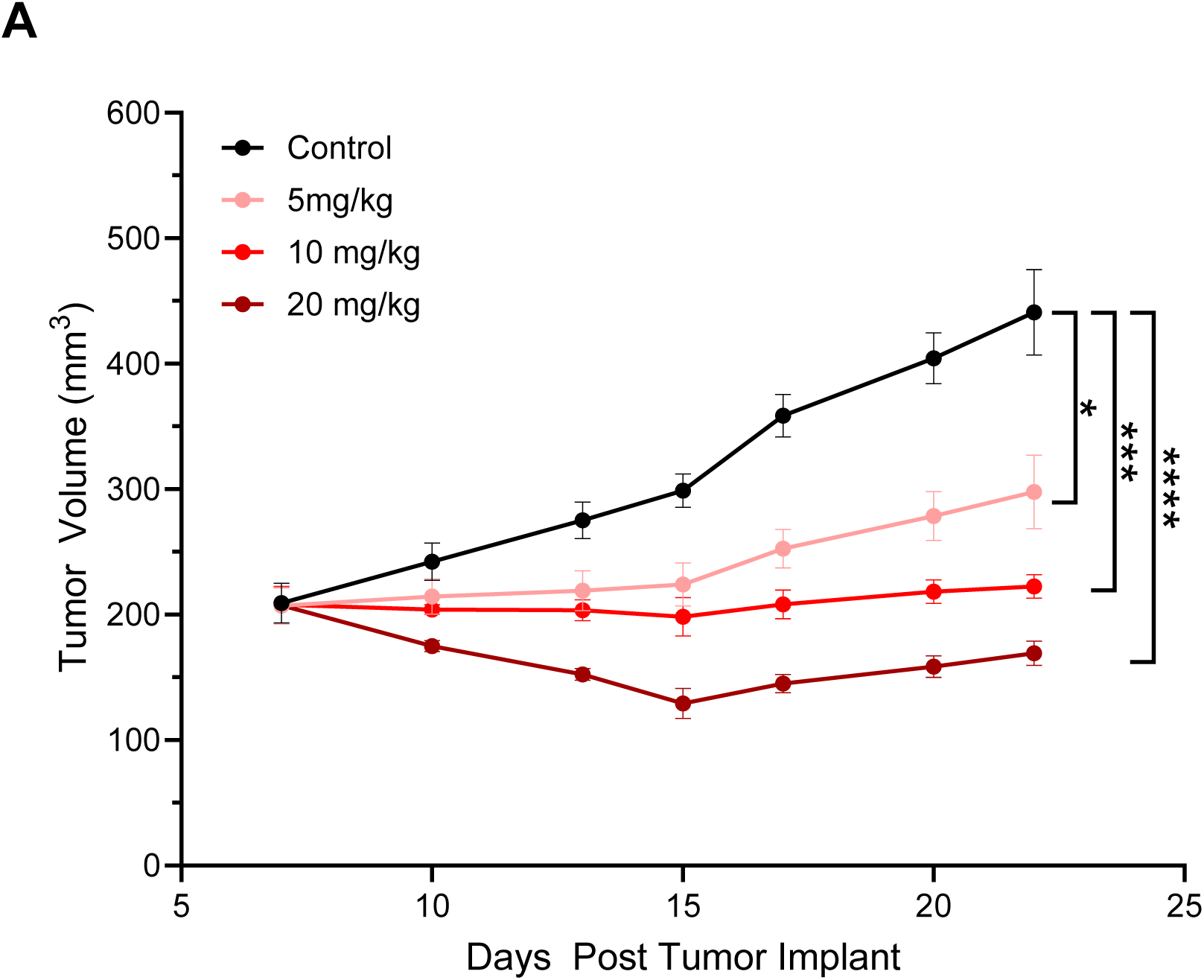
**A**) Tumor volumes for U251 xenotransplants treated with control (vehicle, black) or with the indicated osimertinib dose administered QD (5 mg/kg, pink; 10 mg/kg, red; 20 mg/kg, scarlet). Statistics, 1-way ANOVA (****p<0.0001, ***,p<0.001; *,p<0.05; n=5/group) for volumes at Day 22. Error bars represent standard deviations.

**Supplementary Figure 2.**
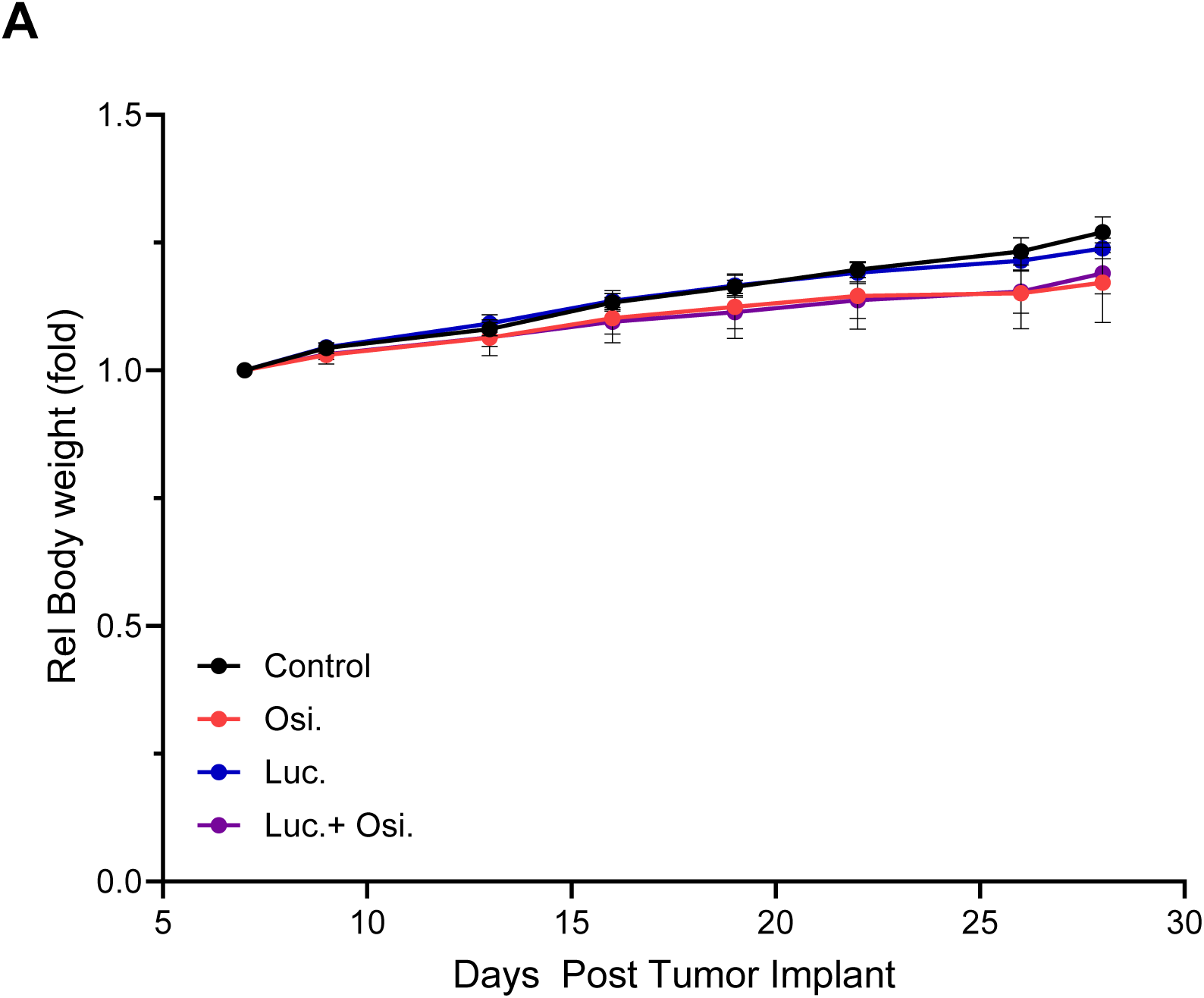
**A**) Relative mouse body weights reported for U251 xenotransplants treated with control (vehicle, black) or with the indicated treatment (subpharmacologic osimertinib, 2 mg/kg QD, red; subpharmacologic lucicebtide 10 mg/kg 3x/week, blue; combination, purple). No statistical significance was found by 1-way ANOVA (***,p<0.001;**,P<0.01; *,p<0.05; ns, not significant; n=5/group) for volumes at Day 30. Error bars represent standard deviations.

